# LplA-mediated rescue of lethal myopathy and cardiomyopathy in mitochondrial lipoylation-deficient mammalian caused by *Lipt1*

**DOI:** 10.64898/2026.11.01.761485

**Authors:** Junru Yu, Zhijuan Hu, Ziwei Liu, Runqing Yang, An-Ping Zeng

**Affiliations:** Center of Synthetic Biology and Integrated Bioengineering, Westlake University; Hangzhou, Zhejiang, China; School of Engineering, Westlake University; Hangzhou, Zhejiang, China; School of Life Sciences, Westlake University; Hangzhou, Zhejiang, China; School of Future Biomedicine, Westlake University; Hangzhou, Zhejiang, China; Research Center for Industries of the Future, Westlake University; Hangzhou, Zhejiang, China; Key Laboratory of Intelligent Low-Carbon Biosynthesis of Zhejiang Province, Westlake University; Hangzhou, Zhejiang, China

**Keywords:** Mitochondrial metabolism, lipoylation deficiency, pathway substitution, lipoate protein ligase, AAV gene therapy, skeletal myopathy, cardiomyopathy

## Abstract

Inherited mitochondrial metabolic disorders impair cellular energy generation and often lead to fatal metabolic crises. Mitochondrial lipoylation defects represent a severe case of such disorders that can lead to lethality but lacks effective treatment so far. In this study, we generated a viable mouse model with muscle-specific deficiency of the lipoyltransferase coded by *Lipt1* to provide an *in vivo* preclinical model that overcomes embryonic lethality. Using this model, we examined a bacterial lipoate protein ligase A (LplA)-mediated gene therapy via adeno-associated virus (AAV) delivery. By introducing LplA alongside lipoic acid supplementation, we reconstituted a salvage pathway that bypasses the mammalian endogenous lipoylation pathway. The intervention rescued premature lethality, recovered motor function and reversed hypertrophic cardiomyopathy in the *Lipt1*-deficient mouse model. Mechanistically, AAV-LplA reconstitutes mitochondrial protein lipoylation and oxidative phosphorylation *in situ*. This energetic rescue suppresses phosphoinositide 3-kinase (PI3K)-Akt hyperactivation, thereby restoring autophagic clearance and subsequently contributing to the recovery of the pathological phenotype. This study established AAV-LplA as a promising gene-delivery strategy for rescuing lethal mitochondrial lipoylation disorders.

## INTRODUCTION

Inherited metabolic disorders (IMDs) that impair mitochondrial energy metabolism are among the most severe genetic diseases and frequently cause early-onset neuromuscular failure and multi-organ dysfunction.^1^ Defective mitochondrial lipoylation is particularly devastating, as lipoylation is required for five vital enzyme complexes of one-carbon and energy metabolisms, namely the glycine cleavage system (GCS) and four 2-oxoacid dehydrogenases including pyruvate dehydrogenase (PDH), 2-oxoglutarate dehydrogenase (OGDH), branched-chain 2-oxoacid dehydrogenase (BCKDH), and 2-oxoadipate dehydrogenase (OADH).^2–4^ These enzymes determine central metabolic processes, coupling oxidative decarboxylation reactions to the tricarboxylic acid (TCA) cycle and enabling amino acids catabolism. Thus, lipoylation defects disrupt both mitochondrial energy production and intermediary metabolism. Human lipoylation depends on a coordinated pathway involving LIPT2 (octanoyltransferase), LIAS (lipoate synthase), and LIPT1 (lipoyltransferase).^5^ Deficiency at any of them can lead to lipoylation defects and subsequent multisystem disturbances. Notably, *LIPT1* deficiency is especially severe, causing hypotonia and fatal systemic metabolic crises, due to its role in transferring the lipoyl groups to target enzymes.^6–10^ No effective treatment is currently available. Dietary lipoic acid and ketogenic diets are ineffective because exogenous lipoic acid cannot be conjugated without the transfer machinery and ketogenic diets remain limited by OGDH inactivity, leaving lipoylation disorders without a mechanism-based treatment.

To overcome this limitation, we established a single-gene bypass strategy based on lipoate protein ligase A (LplA) from *Escherichia coli*.^11^ Unlike the multi-step human pathway, LplA functions as an autonomous system directly ligating exogenous lipoate to mitochondrial target proteins.^12–15^ This concept was also explored in mammalian cell models.^16^ Our previous work showed that mitochondrial-targeted LplA restored function in multiple human cell models, including *LIPT1*, *LIAS*, *LIPT2*, *MECR* or *FDX1* deficiency,^11^ supporting its potential as a unified single-gene intervention for diverse lipoylation genotypes. In a germline knock-in mouse model, mitochondria-targeted LplA rescued the embryonic lethality of the *Lipt1*^-/-^ mice and yielded viable offspring with restored metabolic profiles.^11^ Both germline expression and systemic adeno-associated virus (AAV)-mediated *LplA* delivery were well tolerated,^11,17^ identifying LplA as a promising treatment for genetically diverse lipoylation defects.

Following the conceptual validation of LplA-mediated rescue, the next critical step is to assess its *in vivo* therapeutic potential. Our previous work showed that homozygous lipoylation deficiency causes embryonic lethality, whereas *LIPT1* heterozygotes lack overt phenotypes. Thus, conventional knockout models fail to provide a suitable platform for *in vivo* validation. The previous germline modification is not applicable for patient treatment, as clinical translation requires postnatal, somatic gene delivery. To bridge this gap, a targeted gene delivery strategy must be validated in a primary tissue driving systemic metabolic deficiency. Although *LIPT1* deficiency is multisystemic, skeletal muscle dysfunction with hypotonia and cardiac involvement are common manifestations,^6–10^ making striated muscle an appropriate target for preclinical validation. Furthermore, addressing this major organ-specific burden requires a highly efficient and targeted gene delivery system.

Here, we generated a muscle-specific *Lipt1* knockout mouse that bypasses embryonic lethality. AAV 9 was used for its strong cardiac and skeletal muscle tropism as well as sustained transgene expression.^18–20^ A muscle-specific promoter was incorporated to restrict mitochondria-targeted LplA expression to these tissues.^21^ We demonstrated that AAV-LplA delivery combined with lipoic acid supplementation reconstituted the bacterial salvage pathway *in situ*, alleviated the molecular and functional deficits, supporting AAV-LplA as a promising therapeutic candidate for lethal mitochondrial lipoylation disorders.

## RESULTS

### Intramuscular AAV-LplA gene delivery rescues muscle pathology and motor function in focal *Lipt1*-knockout mice

To quickly evaluate the efficacy of AAV strategy, a tibialis anterior (TA)-specific *Lipt1*-deficient mouse model (Δ*Lipt1*) was constructed at 6 weeks of age by intramuscular injection of AAV-MHCK7-SaCas9-sgRNA (AAV-SaCas9-sgRNA) (Figure 1A and Figure S1A). At 10 weeks of age, the AAV9-MHCK7-MTS-LplA-EGFP (AAV-LplA), which was designed with a muscle-specific chimeric myosin heavy chain and creatine kinase (MHCK7) promoter and a mitochondrial targeting sequence (MTS) to achieve LplA expression in muscle mitochondria, was applied for the sequential rescue by intramuscular administration (Figure 1A). Exogenous lipoic acid was supplemented in drinking water for this strategy, because free lipoic acid released by delipoylation *in vivo* cannot meet the substrate demand of the LplA-dependent salvage pathway (Figure 1A).^11^ *In vivo* imaging of EGFP signals and immunoblotting confirmed LplA expression in the injected muscle (Figure 1B and Figure S1B).

**Figure 1.**
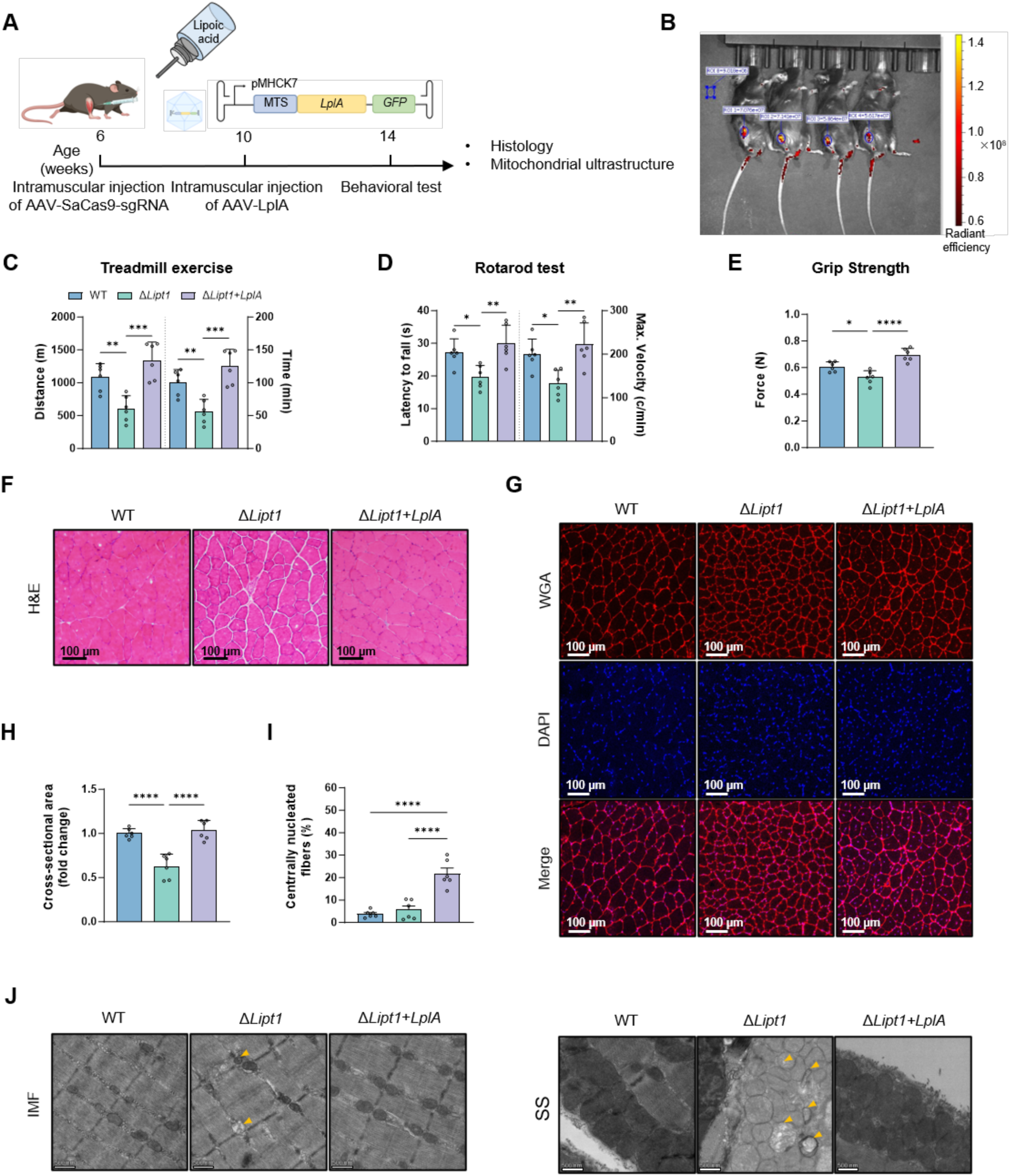
Intramuscular AAV-LplA gene delivery rescues muscle pathology and motor function in a focal model of *Lipt1* deficiency. **(A)** Experimental scheme. AAV-SaCas9-sgRNA was injected into the tibialis anterior (TA) muscle of 6-week-old mice to induce focal *Lipt1* deletion. The same TA muscle was injected with AAV-LplA at 10 weeks of age, and lipoic acid (LA) was supplied in the drinking water. Behavioral tests were performed at 14 weeks of age, followed by tissue collection for histological and ultrastructural analyses. **(B)** Representative *in vivo* fluorescence images after focal TA injection. The EGFP signal is confined to the injected muscle. **(C)** Treadmill running distance (left axis) and running time (right axis) in treadmill exercise test across wild-type (WT), *Lipt*1-deficient (Δ*Lipt1*) and AAV-LplA-treated (Δ*Lipt1*+*LplA*) mice. **(D)** Latency to fall (left axis) and maximum velocity (right axis) in the accelerating rotarod test. **(E)** Maximal hindlimb grip force measured by grip strength test. **(F)** Representative TA cross-sections stained with hematoxylin and eosin (H&E). **(G)** Representative TA cross-sections stained with wheat germ agglutinin (WGA, red) and 4′,6-diamidino-2-phenylindole (DAPI, blue). **(H)** Myofiber cross-sectional area, expressed as fold change relative to WT. **(I)** Percentage of centrally nucleated myofibers. **(J)** Representative transmission electron micrographs of intermyofibrillar (IMF) and subsarcolemmal (SS) mitochondria in TA myofibers. Yellow arrowheads indicate structurally abnormal mitochondria. Scale bars: 100 µm (F and G), 500 nm (J). n = 6 mice per group. Data are presented as mean ± SD. Statistical significance was determined by one-way ANOVA followed by Tukey’s multiple comparisons test. \**P* < 0.05, \*\**P* < 0.01, \*\*\**P* < 0.001 and \*\*\*\**P* < 0.0001.

Given that mitochondrial dysfunction frequently causes muscle weakness and exercise intolerance,^22–24^ we next assessed the functional impact of localized *Lipt1* gene loss and whether AAV-LplA could alleviate resulting deficits. The mice were subjected to standardized functional evaluations at 4 weeks post-treatment (Figures 1C-1E). The *Lipt1*-deficient mice developed exercise intolerance and muscle weakness, with reduced endurance on a forced treadmill (Figure 1C), impaired motor coordination on the rotarod (Figure 1D), and decreased hindlimb grip force (Figure 1E). Intramuscular AAV-LplA administration restored these functional parameters to control levels (Figures 1C-1E). These functional assessments show that acute somatic *Lipt1* deficiency compromises skeletal muscle performance and that local LplA complementation reverses this deficit.

Consistent with the observed functional rescue, local AAV-LplA administration ameliorated the histopathological defects in TA muscles (Figures 1F-1I). Histological examination by hematoxylin and eosin (H&E) and wheat germ agglutinin (WGA) revealed myopathic alterations in untreated *Lipt1*-deficient mice, with myofiber atrophy and enlarged interstitial spaces indicative of early edema or interstitial expansion (Figures 1F and 1G). Morphometric analysis revealed a marked reduction in mean myofiber cross-sectional area in *Lipt1*-deficient TA muscles (Figure 1H). In untreated *Lipt1-*deficient muscles, severe bioenergetic exhaustion compromised endogenous myogenic repair, resulting in defective muscle regeneration and persistent tissue damage. AAV-LplA delivery reversed myofiber atrophy and restored mean myofiber cross-sectional area to physiological dimensions (Figures 1G and 1H). Besides, AAV-LplA treatment also increased the proportion of centrally nucleated myofibers in the injured TA muscle, indicative of new myofiber formation and ongoing muscle repair and regeneration (Figure 1I).^25–28^

The organellar ultrastructure was then examined to determine whether *Lipt1* deficiency perturbed mitochondrial architecture. *Lipt1-*deficient myofibers showed ultrastructural abnormalities, including decreased matrix electron density and inner membrane disruption in both intermyofibrillar (IMF) and subsarcolemmal (SS) mitochondria (Figure 1J). AAV-LplA treatment restored mitochondrial ultrastructural integrity, recovering regular cristae architecture and alignment along the sarcomere lattice (Figure 1J). Restoration of cristae density and sarcomere alignment shows that the structural organization supporting contraction recovers in parallel with muscle function.

### Early AAV-LplA intervention prevents lethality and reverses biochemical abnormalities in *Lipt1*-cKO mice

Following the functional rescue observed in focal muscle, systemic intervention was designed to address the widespread striated muscle and cardiac involvement in clinical mitochondrial myopathies. To facilitate this systemic approach, a striated muscle-specific conditional knockout mouse model (*Ckmm*-Cre; *Lipt1*^flox/flox^, *Lipt1*-cKO) was established using CRISPR-Cas9 technology and the Cre-loxP system (Figure 2A and Figure S1C). These genetically engineered mice successfully bypassed embryonic lethality, thereby providing a viable disease model for subsequent *in vivo* validations. To evaluate systemic intervention, 4-week-old *Lipt1*-cKO mice were intravenously treated with AAV-LplA along with lipoic acid supplementation (Figure 2A). For the assessment of dose-dependent therapeutic efficacy, the mutant mice were divided into distinct groups receiving either a low dose (1 × 10^11^ VG/mouse) or a high dose (1.2 × 10^12^ VG/mouse) of AAV-LplA.

**Figure 2.**
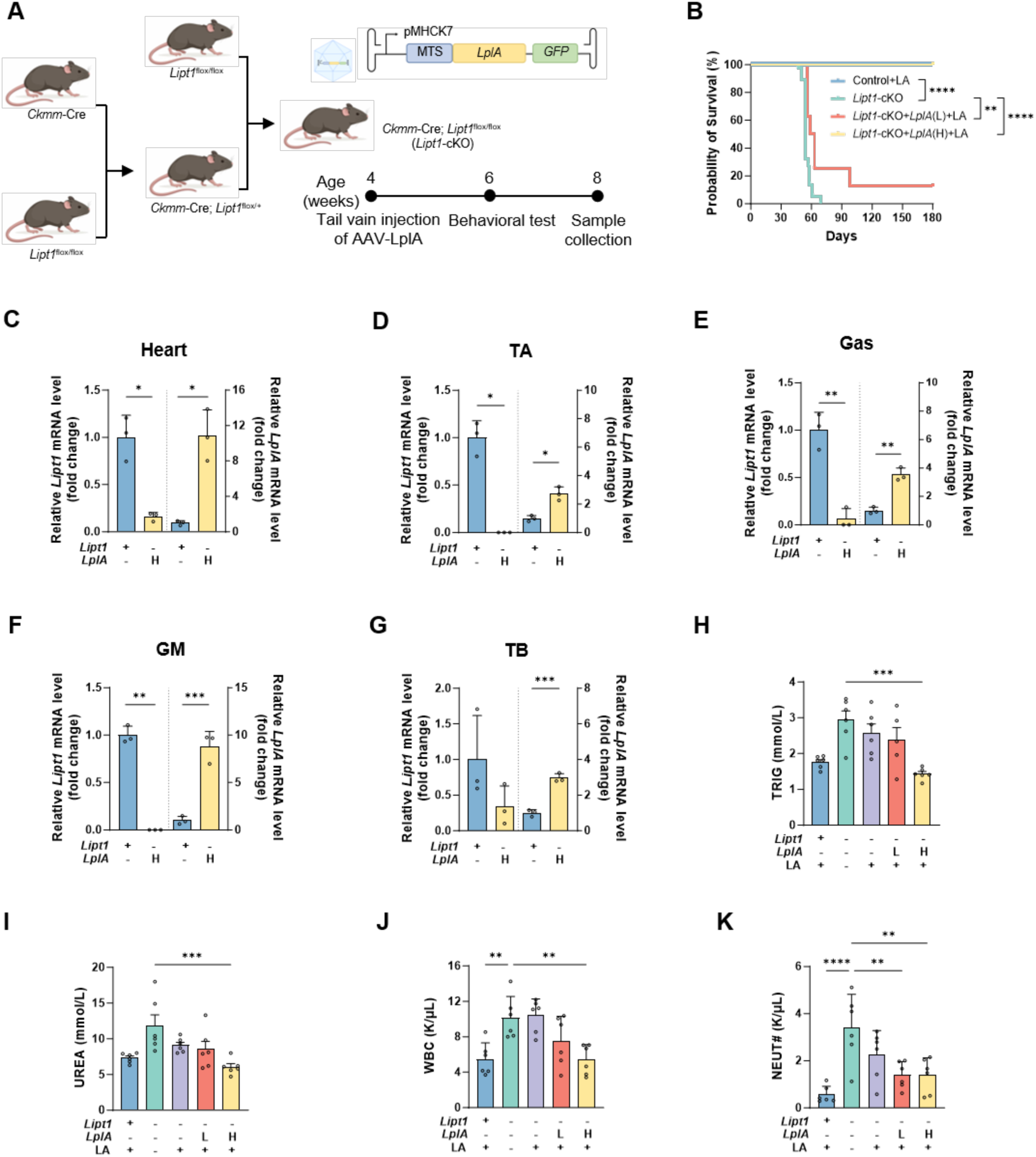
Intravenous AAV-LplA rescues premature lethality and reverses systemic biochemical abnormalities in *Lipt1*-cKO mice. **(A)** Breeding strategy used to generate Ckmm-Cre; *Lipt1*^flox/flox^ (*Lipt1*-cKO) mice and schematic of the experimental timeline. AAV-LplA was delivered by a single tail-vein injection at 4 weeks of age. Behavioral testing was performed at 6 weeks and tissues were collected at 8 weeks of age. **(B)** Survival curves of control mice (*Lipt1*^flox/flox^), *Lipt1*-cKO mice and AAV9-LplA-treated *Lipt1*-cKO mice (*Lipt1*-cKO+LplA) across different vector doses (L, low dose; H, high dose) with or without oral LA supplementation in drinking water. **(C-G)** Relative *Lipt1* (left axis) and *LplA* (right axis) mRNA levels in heart (C), TA (D), gastrocnemius (Gas, E), gluteus maximus (GM, F) and triceps brachii (TB, G), determined by RT-qPCR. *GAPDH* was used as the internal control for normalization. **(H-K)** Plasma triglyceride (TRIG, H), urea (UREA, I), white blood cell count (WBC, J) and neutrophil count (NEUT#, K) in the indicated groups. n = 3 mice per group (C-G), n = 5-6 mice per group (H-K). Survival curves were compared with the log-rank test in B. Statistical significance was determined by unpaired two-tailed t-test in C-G, statistical significance was determined by One-way ANOVA followed by Tukey’s multiple comparisons test in H-K. Data are presented as mean ± SD. \**P* < 0.05, \*\**P* < 0.01, \*\*\**P* < 0.001 and \*\*\*\**P* < 0.0001.

To assess the therapeutic efficacy, longitudinal survival monitoring was performed. Muscle-specific knockout of *Lipt1* induced premature lethality beginning at 47 days of age, whereas littermates with other genotypes showed normal survival (Figure 2B). Systemic AAV-LplA delivery conferred a dose-dependent survival benefit. Low-dose treatment prolonged survival beyond 56 days, and the high-dose delivery prevented premature lethality, with all mice in this group remaining alive and monitoring ongoing (Figure 2B). Reverse transcription quantitative PCR (RT-qPCR) analysis provided the molecular basis for these survival outcomes. Endogenous *Lipt1* expression was nearly abolished across cardiac and diverse skeletal muscles (Figures 2C-2G and Figures S1D-S1G). Minor residual transcripts likely originated from untargeted non-myocyte populations within the tissue homogenates. Exogenous *LplA* transcripts were detected in both cardiac and skeletal muscles of AAV-LplA-treated mice (Figures 2C-2G and Figures S1D-S1G). These results indicated that LplA expression effectively rescued the lethal phenotype of *Lipt1* deficiency.

Human *LIPT1* deficiency is frequently accompanied by systemic metabolic crises.^6–10^ Following the establishment of the viable mouse model, hematological and biochemical analyses were conducted. Untreated *Lipt1*-cKO mice showed elevated serum triglycerides (TRIG) and blood urea (UREA) levels, indicating widespread metabolic derangement and secondary renal stress (Figures 2H and 2I). Dietary lipoic acid supplementation alone did not improve these severe biochemical abnormalities, whereas systemic AAV-LplA therapy reversed these metabolic defects in a dose-dependent manner. While low-dose intervention was insufficient, high-dose systemic AAV-LplA administration restored both TRIG and UREA to baseline levels (Figures 2H and 2I). Circulating white blood cells (WBC) and neutrophils (NEUT) were also increased in untreated *Lipt1*-cKO mice, reflecting secondary systemic inflammation (Figures 2J and 2K). Consistent with the biochemical rescue, high-dose AAV-LplA treatment reversed this leukocytosis and neutrophilia, indicating the resolution of secondary systemic inflammation (Figures 2K and 2L). In addition, intravenous AAV delivery was well tolerated *in vivo*. Erythroid indices, platelet counts, and liver function panels remained well within baseline ranges across all treated mice (Figure S2).

### Systemic AAV-LplA gene delivery preserves myofiber integrity and restores motor performance

Multiple muscles were first surveyed to determine whether systemic AAV-LplA gene delivery could broadly rescue skeletal muscle pathology (Figure 3A). Body weight remained comparable across all experimental groups, but untreated *Lipt1*-cKO mice exhibited variable muscle mass loss (Figure 3B and Figures S3A-S3B). Marked reductions were observed in tibialis anterior (TA), triceps brachii (TB), gastrocnemius (Gas), and gluteus maximus (GM), whereas the extensor digitorum longus (EDL), soleus (Sol), pectoralis major (PM), and quadriceps (QA) showed no detectable change (Figure S3B). Dietary lipoic acid supplementation alone did not significantly attenuate muscle mass loss (Figure 3B). In contrast, intravenous delivery of AAV-LplA dose-dependently rescued these mass deficits, with the high dose restoring relative muscle weights to control levels (Figure 3B). The graded dose response suggests that a threshold of LplA activity is required for muscle preservation.

**Figure 3.**
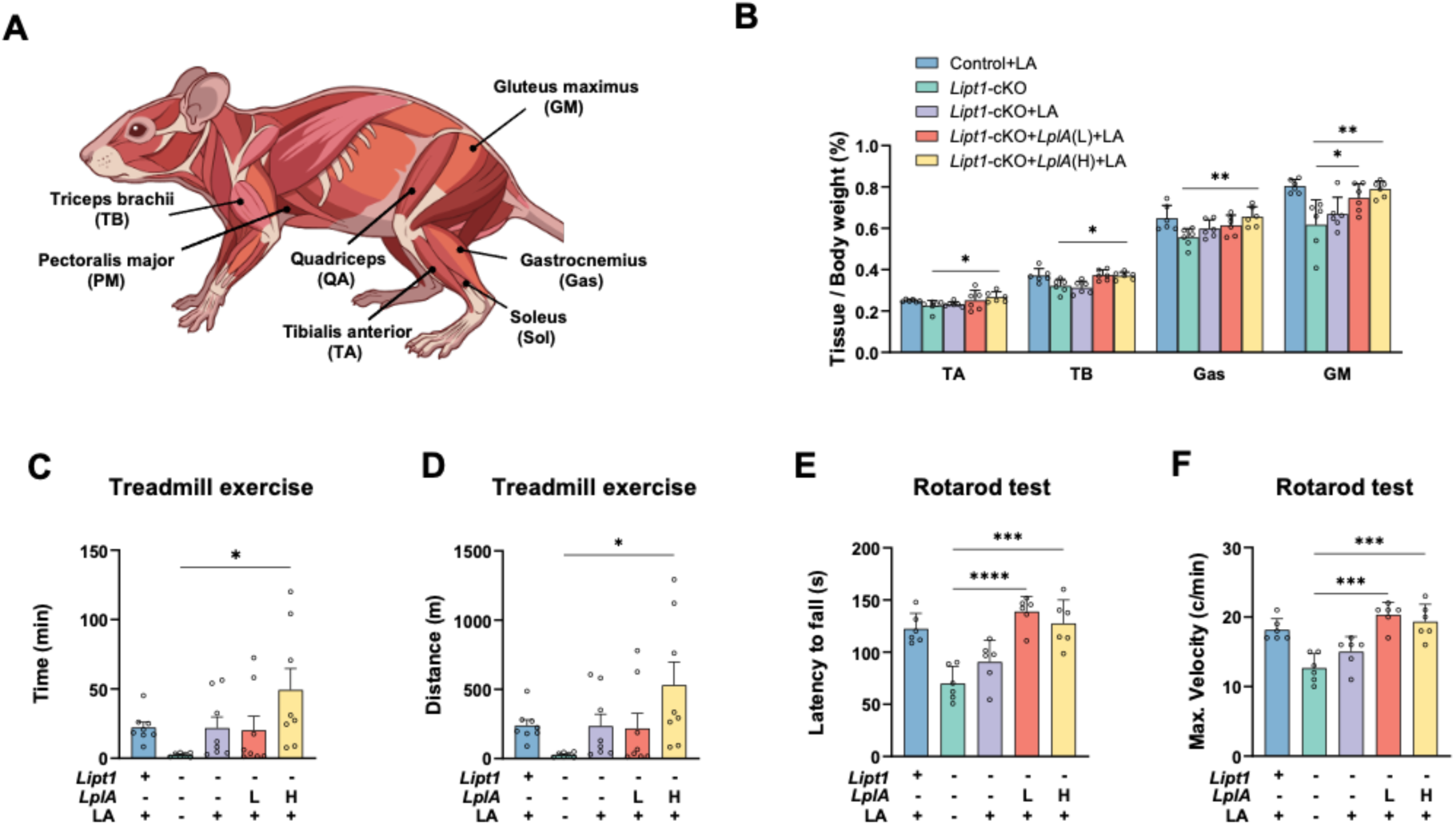
Systemic AAV-LplA restores muscle mass and motor performance in *Lipt1*-cKO mice. **(A)** Schematic of the muscles analyzed, including pectoralis major (PM), quadriceps (QA), TA, extensor digitorum longus (EDL), soleus (Sol), Gas, TB and GM. **(B)** Relative weights of TA, TB, Gas and GM normalized to body weight in Control+LA, *Lipt1*-cKO, *Lipt1*-cKO+LA, *Lipt1*-cKO+*LplA*(L)+LA and *Lipt1*-cKO+*LplA*(H)+LA mice. **(C and D)** Treadmill running time (C) and running distance (D) in the treadmill exercise test. **(E and F)** Rotarod performance, shown as latency to fall (E) and maximum velocity (F). n = 6-8 mice per group. Data are presented as mean ± SD. Statistical significance was determined by one-way ANOVA followed by Tukey’s multiple comparisons test. \**P* < 0.05, \*\**P* < 0.01, \*\*\**P* < 0.001 and \*\*\*\**P* < 0.0001.

We next tested whether these reductions in muscle mass impaired exercise performance. AAV-LplA administration improved the motor deficits observed in untreated *Lipt1*-cKO mice (Figures 3C-3F and Figure S3C). Systemic gene delivery overcame their exercise intolerance, with dose-dependent increases in treadmill running distance and running time (Figures 3C and 3D). It also restored motor coordination and balance, as reflected by improvements in rotarod performance (Figures 3E and 3F). Notably, mice receiving lipoic acid supplementation alone showed a slight upward trend in motor performance, but these changes did not reach statistical significance (Figures 3C-3F). This marginal effect likely reflects the antioxidant capacity of lipoic acid, which can provide nonspecific protection against oxidative stress without correcting the underlying lipoylation defect.^29–31^

To identify the structural basis of impaired motor function, we next performed histological and ultrastructural analyses on the muscles with altered mass (TA, Gas, TB and GM). TA and Gas were prioritized because they are major contributors to hindlimb support and propulsion during locomotion. TB and GM were also included to verify the systemic extent of the pathology and therapeutic rescue. WGA staining showed that AAV-LplA treatment reversed myofiber atrophy in the TA, Gas, TB and GM muscles, restoring cross-sectional area to control levels, whereas lipoic acid supplementation alone conferred no protection against myofiber atrophy (Figures 4A-4C and Figures S3D-S3F). Masson’s trichrome staining revealed collagen deposition and interstitial fibrosis in *Lipt1*-deficient muscles, which were cleared by AAV-LplA administration but remained unaffected by lipoic acid supplementation (Figures 4D-4F and Figures S3G-S3I).

**Figure 4.**
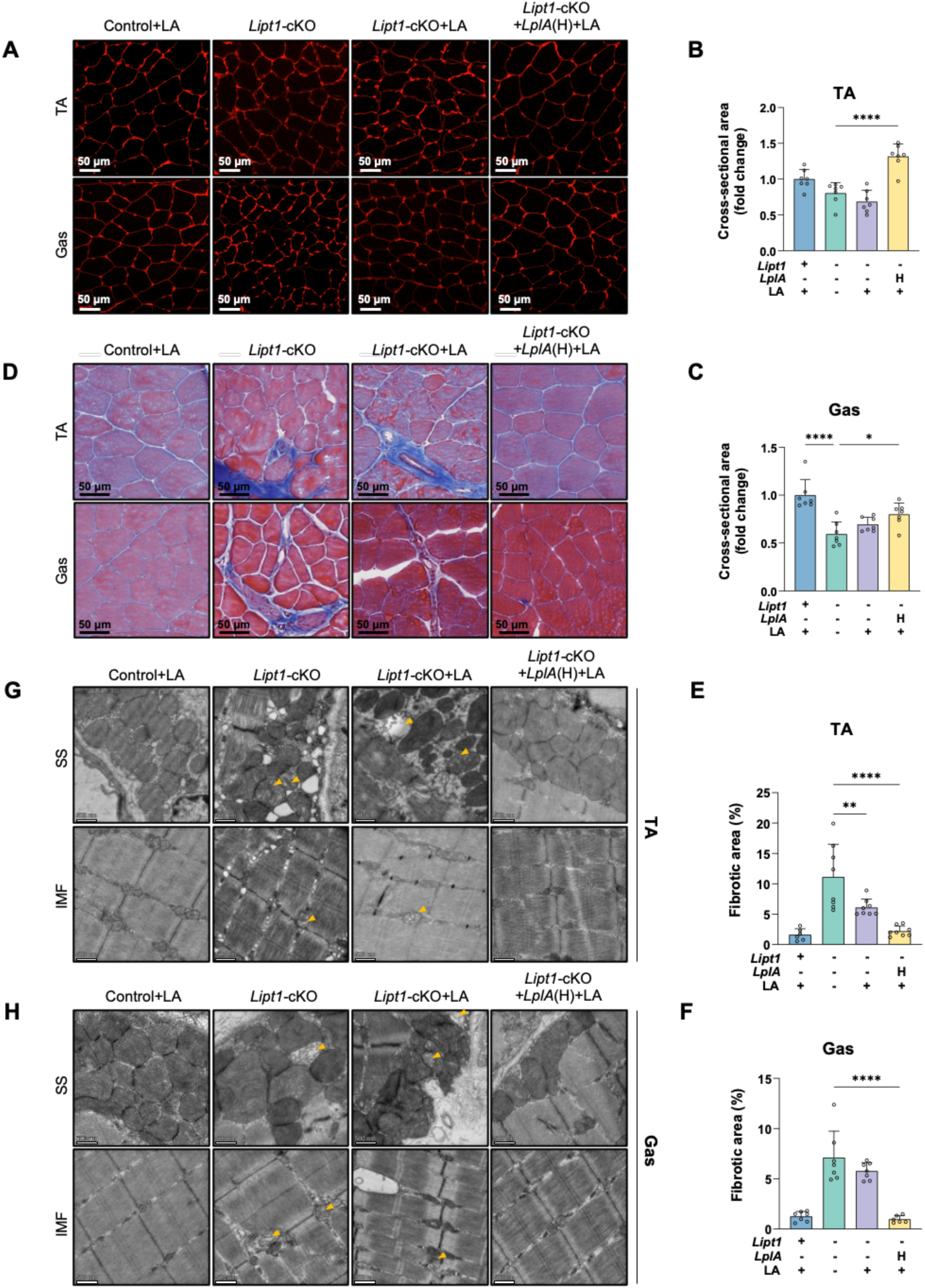
AAV-LplA reverses myofiber atrophy, interstitial fibrosis and mitochondrial ultrastructural defects in *Lipt1*-deficient skeletal muscle. **(A)** Representative WGA-stained cross-sections of TA (top) and Gas (bottom) muscles from Control+LA, *Lipt1*-cKO, *Lipt1*-cKO+LA and *Lipt1*-cKO+LplA(H)+LA mice. **(B and C)** Myofiber cross-sectional area in TA (B) and Gas (C), expressed relative to Control+LA. **(D)** Representative Masson’s trichrome-stained cross-sections of TA and Gas muscles. **(E and F)** Interstitial fibrotic area in TA (E) and Gas (F). **(G and H)** Representative transmission electron micrographs of SS and IMF mitochondria in TA (G) and Gas (H) myofibers. Yellow arrowheads indicate structurally abnormal mitochondria. Scale bars, 50 µm (A and D) and 500 nm (G and H). n = 6-8 mice per group (B, C, E and F). Data are presented as mean ± SD. Statistical significance was determined by one-way ANOVA followed by Tukey’s multiple comparisons test. \**P* < 0.05, \*\**P* < 0.01 and \*\*\*\**P* < 0.0001

To examine the subcellular structures of these affected muscles, transmission electron microscopy (TEM) was used to assess mitochondrial ultrastructure (Figures 4G and 4H). In *Lipt1*-cKO muscles, regardless of lipoic acid supplementation, both IMF and SS mitochondria showed severe morphological defects, including mitochondrial swelling, reduced matrix density and fragmented cristae (Figures 4G and 4H). AAV-LplA treatment corrected these structural abnormalities, restoring dense matrix appearance, regular cristae arrays, and alignment along the sarcomeres (Figures 4G and 4H). Collectively, these data show that systemic LplA reconstitution corrected the structural deterioration of skeletal muscles at both histological and subcellular levels, consistent with the recovery of motor function.

### AAV-LplA ameliorates cardiac hypertrophy and attenuates renal injury

Human lipoylation disorders are frequently present with cardiomyopathy,^7,32,33^ and *Lipt1*-cKO mice developed a similar cardiac phenotype. Relative tissue weight measurements revealed increased heart weight and a reduction in kidney weight in untreated *Lipt1*-cKO mice, indicating cardiac hypertrophy accompanied by secondary renal injury (Figure 5A). Low-dose AAV-LplA intervention produced no statistically significant changes in these organ weights. At the high dose, systemic AAV-LplA delivery restored both heart and kidney masses to control levels, whereas weights of other major non-target organs remained unaltered across all groups (Figure 5A). Therefore, the high-dose group was used exclusively for subsequent structural and molecular evaluations. Hearts from both untreated and lipoic acid-supplemented *Lipt1*-cKO mice showed ventricular wall thickening and chamber dilation. Intravenous AAV-LplA administration reversed this pathological cardiac remodeling (Figure 5B).

**Figure 5.**
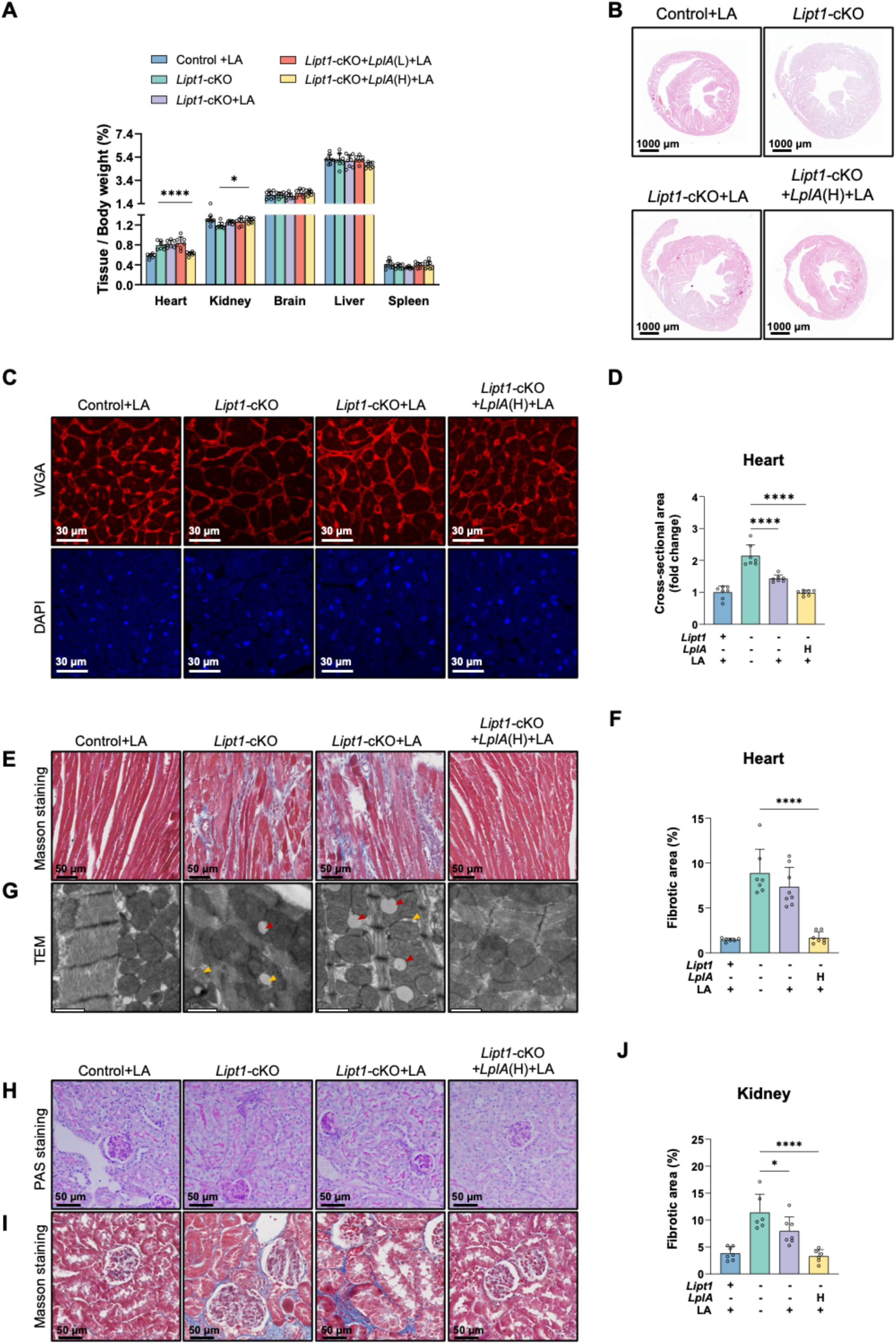
Systemic AAV-LplA ameliorates cardiac hypertrophy and alleviates secondary renal injury in *Lipt1*-cKO mice. **(A)** Weights of heart, kidney, brain, liver and spleen normalized to body weight in Control+LA, *Lipt1*-cKO, *Lipt1*-cKO+LA, *Lipt1*-cKO+*LplA*(L)+LA and *Lipt1*-cKO+*LplA*(H)+LA mice. **(B)** Representative heart sections stained with H&E. **(C)** Representative heart sections stained with WGA (red) and DAPI (blue). **(D)** Cardiomyocyte cross-sectional area, expressed as fold change relative to Control+LA. **(E)** Representative Masson’s trichrome-stained heart sections. **(F)** Quantification of fibrotic area in heart sections. **(G)** Representative transmission electron micrographs of heart mitochondria. Yellow arrowheads indicate structurally abnormal mitochondria. Red arrowheads indicate lipid droplets. **(H)** Representative periodic acid-Schiff (PAS)-stained kidney sections. **(I)** Representative Masson’s trichrome-stained kidney sections. **(J)** Quantification of fibrotic area in kidney sections. Scale bars, 1000 µm (B), 30 µm (C), 50 µm (E, H and I) and 1 µm (G). n = 6-8 mice per group. Data are presented as mean ± SD. Statistical significance was determined by one-way ANOVA followed by Tukey’s multiple comparisons test. \**P* < 0.05 and \*\*\*\**P* < 0.0001.

These changes prompted a more detailed analysis of cardiomyocyte architecture. Consistent with the macroscopic rescue, high-dose AAV-LplA delivery attenuated histological damage and restored cardiomyocyte dimensions in *Lipt1*-cKO myocardium (Figures 5C-5F). WGA staining of cardiac sections showed cardiomyocyte hypertrophy in *Lipt1*-cKO mice, which was marginally reduced by lipoic acid supplementation alone but normalized by AAV-LplA treatment (Figures 5C and 5D). Masson’s trichrome staining revealed interstitial myocardial fibrosis in the hearts of *Lipt1*-cKO and lipoic acid-supplemented mice, which was cleared following AAV-LplA gene transfer (Figures 5E and 5F). At the subcellular level, *Lipt1*-deficient cardiomyocytes showed disrupted mitochondrial integrity with cristae fragmentation (Figure 5G). Intramyocellular lipid droplets also accumulated in *Lipt1*-deficient hearts (Figure 5G). Dietary lipoic acid supplementation did not restore these mitochondrial ultrastructural defects, leaving mitochondrial vacuolation and blurred cristae unchanged. In contrast, these ultrastructural aberrations were reversed following AAV-LplA gene transfer, restoring densely packed cristae arrays throughout the myofibrillar lattice and clearing pathological lipid droplets (Figure 5G).

Prompted by the recovery of kidney weight, we next examined the renal histology. Periodic acid-Schiff (PAS) staining revealed degenerative lesions in the renal cortex of *Lipt1*-cKO mice, featuring glomerulosclerosis and cellular atrophy (Figure 5H). Masson’s trichrome staining demonstrated interstitial collagen deposition and renal fibrosis in *Lipt1*-cKO mice (Figures 5I and 5J). While dietary lipoic acid supplementation conferred limited therapeutic benefit against these renal lesions, systemic AAV-LplA delivery attenuated cortical degeneration and cleared interstitial fibrotic deposition, restoring renal tissue architecture (Figures 5I and 5J). The parallel improvement in cardiac and renal architecture may point to a hemodynamic contribution to the renal lesion.^34^

### AAV-LplA reconstitutes mitochondrial lipoylation and reverses cardiac molecular abnormalities

To define the molecular basis by which AAV-LplA treatment rescues *Lipt1*-deficient mice, we examined the mitochondrial protein lipoylation levels across multiple striated muscles *in vivo*. In untreated *Lipt1*-cKO mice, lipoylation of dihydrolipoamide acetyltransferase (DLAT) and dihydrolipoamide succinyltransferase (DLST) was nearly undetectable in the heart, TA, Gas, TB and GM (Figures 6A-6E). The faint residual signals detected in these muscles likely originated from untargeted non-myocyte populations within tissue isolates (Figures 6A-6E). Dietary lipoic acid supplementation alone failed to improve lipoylation levels (Figures 6A-6E). This failure occurs because mammalian LIPT1 functions exclusively to transfer lipoyl groups and cannot directly use free lipoic acid. Conversely, intravenous AAV-LplA administration restored protein lipoylation across these muscle tissues. Restoration of DLAT and DLST lipoylation across all examined muscles confirms that the bacterial enzyme reconstitutes mitochondrial protein lipoylation *in vivo*. Given the central role of the heart in systemic metabolic homeostasis, we focused subsequent analyses on cardiac tissue and assessed the abundance of oxidative phosphorylation (OXPHOS) complexes, which function downstream of lipoylation-dependent mitochondrial metabolism. *Lipt1* deficiency reduced myocardial OXPHOS subunit levels (Figure 6F), consistent with the requirement for lipoylation in TCA cycle dehydrogenase activity, which supports respiratory chain function. High-dose AAV-LplA restored the expression of multiple OXPHOS subunits (Figure 6F), indicating the recovery of mitochondrial oxidative metabolism in the *Lipt1*-deficient heart.

**Figure 6.**
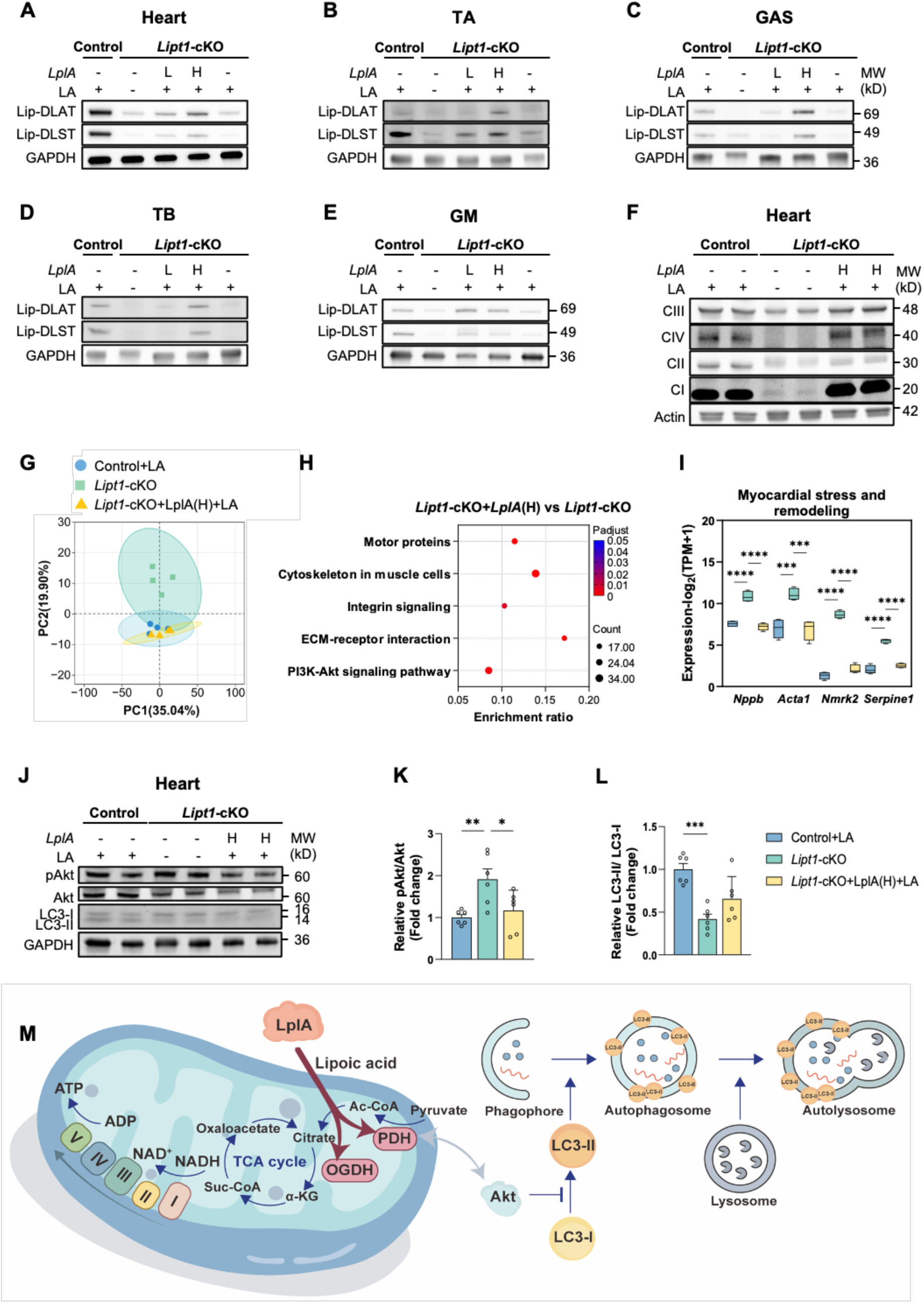
Systemic AAV-LplA reconstitutes mitochondrial metabolism and reverses cardiac molecular defects. **(A-E)** Immunoblots of lipoylated DLAT (Lip-DLAT) and DLST (Lip-DLST) in heart (A), TA (B), Gas (C), TB (D) and GM (E). GAPDH was used as the control. **(F)** Immunoblots of oxidative phosphorylation (OXPHOS) subunits from complexes NDUFB8 (CI), SDHB (CII), UQCRC2 (CIII) and MTCO1 (CIV) in heart. Actin was used as the control. **(G)** Principal component analysis of cardiac transcriptomes from Control+LA, *Lipt1*-cKO and *Lipt1*-cKO+*LplA*(H)+LA mice. **(H)** KEGG pathway enrichment of differentially expressed genes. The color scale represents the adjusted P value, and the dot size represents the number of genes. **(I)** Expression of myocardial stress and remodeling genes (*Nppb*, *Acta1*, *Nmrk2* and *Serpine1*). Transcript abundance is shown as log2(TPM+1), where TPM denotes transcripts per million. **(J)** Immunoblots of phosphorylated Akt (pAkt), total Akt, LC3-I and LC3-II in heart. GAPDH was used as the control. **(K)** Quantification of Akt phosphorylation (pAkt/Akt ratio) in hearts from (J). **(L)** Quantification of LC3-II/LC3-I ratio in hearts from (J). **(M)** Schematic illustration of LplA-mediated rescue in lipoylation-deficient hearts. LplA restores lipoylation of the TCA cycle dehydrogenases PDH and OGDH, sustaining TCA cycle activity and oxidative phosphorylation (OXPHOS). Loss of lipoylation impairs TCA cycle and OXPHOS function and produces a bioenergetic deficit that is proposed to activate PI3K-Akt signaling. Akt restrains LC3-I lipidation to LC3-II, limiting autophagosome formation and autophagosome-lysosome fusion, thereby reducing autophagic capacity and contributing to hypertrophic remodeling. The T-bar indicates inhibition. n = 4 mice per group (G-I), n = 6 mice per group (K and L). Data are presented as mean ± SD. Statistical significance was determined by one-way ANOVA followed by Tukey’s multiple comparisons test. \**P* < 0.05, \*\**P* < 0.01 and \*\*\**P* < 0.001.

To investigate how these bioenergetic alterations contribute to pathological myocardial hypertrophy, we performed RNA sequencing (RNA-seq) on heart tissue (Figures 6G-6I and Figure S4). Principal component analysis separated control from *Lipt1*-cKO samples, whereas AAV-LplA-treated groups clustered closely with control mice, indicating broad restoration of the transcriptome (Figure 6G). Volcano plots showed widespread gene expression changes in the impaired *Lipt1*-cKO heart, including marked upregulation of remodeling genes (Figure S4A). Systemic AAV delivery largely reversed these alterations (Figures S4B-S4C). KEGG pathway enrichment analysis further identified the biological pathways associated with these differentially expressed genes (DEGs) (Figure 6H and Figure S4D). The enriched pathways matched the pathological phenotypes described above, and quantitative profiling showed marked induction of pathological marker genes in the *Lipt1*-deficient myocardium (Figure 6I and Figures S4E-S4F). The upregulation of *Nppb* and *Acta1* reflected cardiac hypertrophy and fetal gene reprogramming, while increased *Nmrk2* and *Serpine1* indicated metabolic decompensation and active profibrotic remodeling (Figure 6I). Systemic AAV-LplA delivery suppressed these transcripts and restored their expression to baseline levels (Figure 6I). Enrichment of extracellular matrix (ECM)-receptor interactions and integrin signaling tightly paralleled the interstitial fibrosis that may contribute to impaired cardiac contractility (Figures S4D and S4E). Transcript levels of the fibrillar collagen genes *Col1a1*, *Col3a1*, *Tnc* and *Ccn1* were increased, consistent with increased fibrotic ECM deposition (Figure S4E). Ultrastructural myofibrillar damage was accompanied by alterations in motor proteins and the muscle cytoskeleton pathways (Figure S4D). Sarcomere and cytoskeletal genes, including *Csrp3*, *Xirp2*, *Nrap*, and *Flnc*, showed marked expression changes in *Lipt1*-deficient hearts (Figure S4F). AAV-LplA treatment restored these profibrotic and cytoskeletal transcripts (Figures S4E and S4F).

In addition, KEGG analysis identified enrichment of the phosphoinositide 3-kinase (PI3K)-Akt pathway in *Lipt1*-deficient hearts (Figure 6H), which was further supported by immunoblotting showing increased protein kinase B (Akt) phosphorylation (Figures 6J and 6K). These findings indicate an association between PI3K-Akt activation and the disease phenotype. In the same deficient hearts, the conversion of cytosolic microtubule-associated protein 1 light chain 3 (LC3)-I to autophagosome membrane-bound LC3-II was markedly suppressed, suggesting reduced autophagosome formation (Figures 6J and 6L). Together, these observations indicate that Akt hyperactivation impair LC3-dependent autophagosome formation and autophagic clearance in *Lipt1*-deficient cardiomyocytes, contributing to hypertrophy (Figure 6M). AAV-LplA treatment normalized Akt phosphorylation and relieved this autophagic impairment. By restoring mitochondrial lipoylation and OXPHOS, AAV-LplA eliminated the metabolic defect that drives Akt hyperactivation and impaired autophagy, thereby reversing pathological cardiac phenotype.

## DISCUSSION

Inherited lipoylation deficiencies caused by pathogenic *LIPT1* variants usually present in early life with systemic bioenergetic failure and are often fatal. Human cells lack an autonomous salvage pathway for free lipoic acid as found in bacteria, so dietary supplementation of lipoic acid alone fails to correct the defect.^15,32^ Our previous work showed that the *E. coli*-derived LplA bypass functions effectively in lipoylation-deficient mammalian cells and germline models.^11^ However, achieving *in vivo* phenotypic rescue via postnatal somatic gene delivery remained unexplored. In this study, we demonstrate that AAV9-mediated delivery of bacterial LplA driven by the muscle-specific MHCK7 promoter reconstitutes mitochondrial lipoylation and converts a lethal phenotype into long-term survival with recovery of skeletal muscle and cardiac pathology in a mouse model.

The coordinated recovery of skeletal muscle and heart pathology supports the therapeutic efficacy of AAV-LplA treatment in mammalian. The treatment restored motor function, accompanied by the restoration of myofiber architecture and alleviation of fibrosis (Figures 1C-J, Figures 3C-F, Figure 4 and Figures 5B-5G). The heart, which is composed primarily of cardiac muscle, relies heavily on mitochondrial metabolism. Cardiac involvement is a recognized clinical feature of lipoylation disorders.^7,32,33^ In the *Lipt1-*deficient model, the untreated mice developed cardiac hypertrophy and chamber dilation. AAV-LplA reversed the remodeling program that sustains hypertrophy, with coordinated recovery of its fibrotic and structural components (Figures 6H-6I and Figures S4D-S4F). Besides, lipid droplet accumulation in *Lipt1*-cKO mice suggests impaired mitochondrial fatty acid oxidation and potential lipotoxicity. AAV-LplA treatment cleared intramyocellular lipid droplets, likely reflecting restored TCA cycle function and β-oxidation in metabolically demanding striated muscles.^35^ Nevertheless, a small subset of transcripts, including circadian clock regulators, remained differentially expressed in AAV-LplA-treated hearts relative to controls, and the functional relevance of this residual signature remains to be determined (Figure S4C).

The therapeutic efficacy of AAV-LplA fundamentally stems from the recovery of mitochondrial energy metabolism. *Lipt1* is required for lipoylation of mitochondrial dehydrogenase complexes. Loss of this modification impairs their catalytic activity and disrupts the TCA cycle.^5^ A recent study identified that defective mitochondrial S-adenosylmethionine (mitoSAM)-dependent lipoylation compromises respiratory chain and metabolic function, leading to early cardiomyopathy.^36^ In our study, we addressed the direct lipoylation defect caused by *Lipt1* loss in striated muscle, in which lipoylation was nearly abolished, thereby lowering OXPHOS subunit expression and compromising mitochondrial oxidative metabolism (Figure 6F). PI3K-Akt signaling pathway, a principal regulator of anabolic metabolism and cell growth,^37,38^ was enriched in *Lipt1*-deficient hearts, and immunoblotting confirmed increased Akt phosphorylation (Figures 6H and 6J-6K). These results suggest that PI3K-Akt signaling is activated in response to metabolic stress. Akt activity suppresses autophagosome initiation.^39,40^ The LC3-II/LC3-I ratio was markedly reduced in the same hearts, indicating decreased autophagosome formation and possibly impaired autophagic clearance (Figures 6J and 6L). These findings support a model in which loss of lipoylation impairs OXPHOS, resulting in a bioenergetic deficit that activates PI3K-Akt signaling, which in turn suppresses LC3-dependent autophagosome formation, reducing autophagic capacity and contributing to hypertrophic remodeling (Figure 6M). AAV-LplA reversed this cascade by correcting the primary lipoylation defect and restoring the metabolic impairment, thereby normalizing Akt phosphorylation and autophagosome formation, ultimately reversing cardiac hypertrophy.

Beyond local skeletal and cardiac muscle repair, AAV-LplA treatment also attenuated abnormalities in the kidney. Untreated *Lipt1*-cKO mice developed kidney atrophy and renal injury, together with elevated blood urea, a marker of renal dysfunction (Figure 2I and Figures 5H-5J). The treatment attenuated cardiac hypertrophy and chamber dilation, followed by reduced renal injury.^41^ We speculate that the renal changes arise from impaired systemic perfusion secondary to cardiac dysfunction, and the reduced renal injury after treatment is associated with the cardiac improvement. Together, these observations suggest that correction of primary cardiac defect may contribute to protection against systemic injury.

This study demonstrates the efficacy of AAV-LplA treatment in a *Lipt1*-deficient model. LplA bypasses the human *de novo* lipoylation pathway by directly ligating free lipoic acid to mitochondrial dehydrogenases. This ability may extend the therapeutic utility of LplA to defects across the human lipoylation pathway and in its regulators. LplA has been shown to restore mitochondrial function in human cells with defects in other lipoylation-related genes, including *LIPT2, LIAS, MECR* and *FDX1.*^11,16^ These studies support the potential of LplA to bypass genetically distinct lipoylation defects with a single transgene, which addresses a central limitation of conventional one-to-one gene replacement. The mammalian lipoylation pathway is highly interdependent, and pathogenic variants in genes such as *LIPT1, LIPT2* and *LIAS* can produce similar downstream bioenergetic failure.^33,42–44^ Developing customized vectors for each rare genetic target presents manufacturing and regulatory challenges. By introducing the bacteria-derived LplA with lipoic acid supplementation, this approach enables a lipoylation pathway substitution that offers a broader conceptual framework for metabolic gene therapy. It shows that repurposing evolutionary enzymes can bypass several defects in human pathways. AAV-LplA treatment provides a unified approach for multiple lipoylation defects, although validation across additional genotypes will be required. Further studies should extend this strategy to other lipoylation-related deficiencies.

Furthermore, targeting a non-mammalian enzyme for therapeutic development requires balancing therapeutic efficacy against systemic toxicity.^20,45^ To limit these risks, we combined the myotropic AAV9 capsid with the MHCK7 promoter.^21^ This dual-targeting design was intended to direct LplA expression primarily to cardiac and skeletal muscles. Consistent with the sustained expression and tolerability of LplA reported previously,^11,17^ expression of the exogenous bacterial enzyme caused no apparent systemic toxicity (Figure S2). Liver transaminases, renal function markers, and systemic inflammatory indices remained comparable to those of healthy controls. However, human *LIPT1* deficiency is associated with multi-system disorders. Future clinical translation will require broader systemic promoters to address other multi-organ manifestations. Moreover, therapeutic durability remains an important question to be further studied in AAV-based gene therapy. AAV vectors can sustain long-term episomal transgene expression in post-mitotic tissues,^18^ but it remains unclear whether a single dose at an early age will provide lifelong protection or require booster dosing if episomal genomes are diluted during postnatal muscle growth.^20^ Long-term immune tolerance to the exogenous bacterial protein also requires extended follow-up.

In summary, this study demonstrates that AAV-LplA gene delivery reconstitutes mitochondrial protein lipoylation and provides a foundation for gene therapies targeting fatal lipoylation diseases. By reconstituting the salvage lipoylation pathway in mammalian striated muscle, our findings support further evaluation of AAV-LplA as a promising therapy for lethal mitochondrial lipoylation disorders.

## MATERIAL AND METHODS

### Mice

All animal experimental procedures were approved by the Institutional Animal Care and Use Committee (IACUC) of Westlake University (AP#23-074-2-ZAP) and were performed in accordance with institutional guidelines. Mice received humane care and were maintained in a specific pathogen-free (SPF) facility under controlled environmental conditions (ambient temperature 22 ± 1°C and relative humidity 40-70%) with a 12-h light/dark cycle. Standard chow and drinking water were provided *ad libitum*. Both male and female mice were used in this study. Male and female mice were included in comparable numbers, with both sexes represented across genotypes and treatment groups, and sex was balanced among experimental groups.

To generate muscle-specific conditional knockout animals, *Lipt1*^flox/flox^ homozygotes (generated by intercrossing *Lipt1*^flox/+^ mice) were first crossed with *Ckmm*-Cre mice to obtain *Ckmm*-Cre; *Lipt1*^flox/+^ progeny. These progenies were subsequently bred with *Lipt1*^flox/flox^ mice to yield *Ckmm*-Cre; *Lipt1*^flox/flox^ offspring (*Lipt1*-cKO). Age- and sex-matched *Lipt1*^flox/flox^ littermates were used as controls in all experiments. Genotyping was performed by PCR using genomic DNA extracted from toe clips with the following primers: Flox-F1 (5’-GGACATTGACCTCTGACCTGT-3’), Flox-R1 (5’-ACATGTCCCCCTGATTGTGT-3’), *Ckmm*-Cre-F1 (5’-GTGAAACAGCATTGCTGTCACTT-3’), *Ckmm*-Cre-R1 (5’-TAAGTCTGAACCCGGTCTGC-3’), *Ckmm*-Cre-F2 (5’-CAAATGTTGCTTGTCTGGTG-3’), and *Ckmm*-Cre-R2 (5’-GTCAGTCGAGTGCACAGTTT-3’).

For lipoic acid supplementation, drinking water containing 0.1% (w/v) lipoic acid (LA, S352645, Aladdin, Shanghai, China) was administered to the following groups: (i) *Lipt1*^flox/flox^ mice (Control+LA), (ii) *Ckmm-*Cre; *Lipt1*^flox/flox^ mice receiving LA alone (*Lipt1-*cKO + LA), and (iii) *Ckmm-*Cre; *Lipt1*^flox/flox^ mice receiving AAV-LplA and LA (*Lipt1-*cKO + AAV-LplA + LA). Untreated *Lipt1*-cKO littermates received regular drinking water and served as the disease control group.

### Systemic administration of AAV-LplA in mice

For systemic administration, recombinant adeno-associated virus serotype 9 (AAV9) vectors encoding MTS-LplA (pAAV-MHCK7-MTS-LplA-EGFP) were purchased from YiXueSheng Biosciences Inc. (Shanghai, China). Viral titers were determined and expressed as vector genomes (VG) per milliliter (VG/mL). Tail-vein injection was administered to the experimental mice at two designated dose levels, comprising a low dose (1 × 10¹¹ VG/mouse) or high dose (1.2 × 10¹² VG/mouse), according to the experimental group assignment.

For localized intramuscular delivery, 6-week-old C57BL/6J mice were anesthetized via isoflurane inhalation, and the targeted hindlimbs were shaved and disinfected. A total volume of 15-25 µL of AAV9-MHCK7-saCas9-sgRNA (5 × 10¹² VG/mL) was delivered into the tibialis anterior (TA) muscle using a microsyringe. To ensure uniform viral distribution, a five-point injection pattern was employed (one central site and four peripheral sites, yielding 3-5 µL per site). Four weeks post-injection, the same TA muscle was re-injected with the second AAV9 vector (pAAV-MHCK7-MTS-LplA-EGFP, 5 × 10¹² VG/mL) following the same five-point injection protocol and volume.

### *In vivo* imaging

Whole-body fluorescence imaging was performed using an IVIS Spectrum small animal imaging system (PerkinElmer, Waltham, MA, USA). Prior to image acquisition, mice were anesthetized via isoflurane inhalation, and the hair over the target regions was carefully removed using electric clippers to minimize light scattering and tissue autofluorescence. The regional EGFP fluorescence intensity and signal distribution were captured and analyzed using Living Image software (PerkinElmer, Waltham, MA, USA).

### Rotarod test

Motor coordination and balance were evaluated using an accelerating rotarod apparatus (Xinruan Information Technology, Shanghai, China). To minimize stress responses to the testing environment, mice were acclimatized to the apparatus for three consecutive days prior to formal testing. During the acclimation phase, mice were placed on the rotating rod at a constant speed of 5 rpm for 5 min. For the formal test, an accelerating protocol was employed, wherein the rotation speed linearly increased from 5 to 40 rpm over a 5-min period. The trial was automatically terminated when the mouse fell off the rod and triggered a bottom pressure sensor. The latency to fall (in seconds) and the maximum speed achieved at the time of falling were recorded. Each mouse was subjected to three consecutive trials with a minimum inter-trial rest interval of 15 min to prevent fatigue. The average values from the three trials were used for subsequent statistical analysis.

### Treadmill exercise test

The exhaustive running test was performed using a treadmill apparatus (Yiyan, Jinan, China). To minimize stress responses to the treadmill environment, mice underwent low-intensity acclimation training for five consecutive days prior to the formal test. The training speeds were set at 10, 12, 14, 16, and 18 m/min on days 1 through 5, respectively, with each session lasting 15 min. The formal test was performed at a constant speed of 18 m/min. A mild electrical stimulus was delivered through a shock grid placed at the rear of each lane. Mice were considered to be exhausted when the mouse’s hindlimbs remained on the electric grid for more than 30 s without resuming running on three separate occasions within a 10-min observation period. At that point, the treadmill was stopped immediately, and the total running time and distance were recorded for each animal.

### Grip strength test

Muscle strength was assessed using a grip strength meter (Calvin, Nanjing, China). Mice were positioned to grasp the metal grid of the transducer and gently pulled backward horizontally by the tail at a steady pace until the grip was released. The maximal force was recorded for each attempt. Three independent trials were performed per animal, and the mean values were used for final statistical evaluation.

### Blood collection and analysis

Blood sampling was performed via puncture of the submandibular vein. Briefly, mice were gently restrained, and a sterile 5-mm lancet was used to puncture the submandibular vein. Blood drops were collected directly into EDTA-K₂-coated microtubes (NPK230505; SANLI Medical, Hunan, China) for hematological evaluation, and into lithium heparin-coated tubes (NPLH0505; SANLI Medical, Hunan, China) for plasma biochemistry.

For the complete blood count (CBC), whole blood collected in EDTA-K₂ tubes (NPK230505, SANLI Medical, Hunan, China) was analyzed directly using a ProCyte Dx® Hematology Analyzer (IDEXX Laboratories, ME, USA).

For biochemical analysis, the lithium heparin tubes were gently inverted to ensure thorough mixing with the anticoagulant, followed by centrifugation at 12,000 × g for 10 min to isolate the plasma fraction. Subsequently, approximately 300 µL of the separated plasma was transferred into a Catalyst® sample cup and analyzed using a Catalyst One® Chemistry Analyzer (IDEXX Laboratories, ME, USA) in accordance with the manufacturer’s protocols.

### Histological analysis

Hearts and kidneys were dissected and fixed in 4% paraformaldehyde (PFA) for 24-48 h at room temperature. Following fixation and 75% ethanol washes, the tissues were dehydrated through a descending graded ethanol series using a HistoCore PEARL tissue processor (Leica Biosystems, Wetzlar, Germany) and embedded in paraffin using an Arcadia-1 embedding station (Leica Biosystems, Wetzlar, Germany). Skeletal muscles were embedded in optimal cutting temperature (OCT) compound, snap-frozen in liquid nitrogen-cooled isopentane, and maintained at −80°C until use.

Paraffin-embedded specimens and OCT-embedded muscle tissues were sectioned at thicknesses of 5 μm and 10 μm, respectively. Both types of sections were subjected to standard hematoxylin and eosin (H&E) staining using an ST5020 Multistainer (Leica Biosystems, Wetzlar, Germany).

To further evaluate extracellular matrix remodeling and fibrosis, adjacent paraffin sections were processed using a periodic acid-Schiff (PAS) staining kit (C0142M, Beyotime Biotechnology, Shanghai, China) and a Masson’s trichrome staining kit (DC0033, Leagene Biotechnology, Beijing, China) in accordance with the manufacturers’ instructions. All histological slides were fully digitized using a ZEISS Axioscan 7 digital slide scanner (Carl Zeiss AG, Oberkochen, Germany).

### Transmission electron microscopy

Freshly dissected tissue samples were immediately immersed in a fixative containing 2% PFA and 2.5% glutaraldehyde (GA) in 0.1 M sodium cacodylate buffer (pH 7.2) and incubated overnight at 4 °C. Following three consecutive washes in the 0.1 M sodium cacodylate buffer, the samples were post-fixed with 1% osmium tetroxide in the same buffer for 1 h at 4 °C. The tissues were then rinsed three times with the buffer, followed by three washes in distilled water, and subsequently block-stained with 1% aqueous uranyl acetate for 1 h at ambient temperature. Dehydration was accomplished using an ascending ethanol gradient (30%, 50%, 70%, and 95% for 10 min each), culminating in three 10-min changes of absolute ethanol, followed by a final transition step in pure acetone. For tissue embedding, the specimens were sequentially infiltrated with acetone and EPON 812 resin mixtures prepared at volume ratios of 2:1 and 1:2 for 30 min each. Final infiltration was performed under vacuum using pure EPON 812 resin, with three fresh resin exchanges every 3 h, and allowed to proceed overnight at room temperature. Polymerization was conducted in an oven at 60 °C for 24-48 h. Ultrathin sections (∼70 nm) were generated using an ultramicrotome and mounted onto standard copper grids. Ultrastructural examination of mitochondrial morphology and myofibrillar architecture was conducted using a Talos L120C G2 transmission electron microscope (Thermo Fisher Scientific, MA, USA) operating at an accelerating voltage of 120 kV.

### Wheat germ agglutinin (WGA) staining

Skeletal muscle specimens were processed into either 10-µm cryosections or 5-µm paraffin-embedded sections. Cryosections were fixed with 4% paraformaldehyde (PFA) for 10 min at room temperature, followed by three 5-min rinses in Hanks’ Balanced Salt Solution (HBSS).

Paraffin sections underwent standard deparaffinization in xylene and rehydration through a descending ethanol series. Heat-induced epitope retrieval was subsequently performed by microwaving the slides in a citrate antigen retrieval solution (P0081, Beyotime Biotechnology, Shanghai, China) for 15 min. Following cooling and thorough washing in phosphate-buffered saline (PBS), all sections were subjected to an identical downstream staining protocol.

Tissue sections were incubated in the dark for 15 min at room temperature with Alexa Fluor™-conjugated WGA (1:200 dilution in HBSS; Thermo Fisher Scientific, MA, USA). After three subsequent washes with HBSS, the slides were coverslipped using a 4′,6-diamidino-2-phenylindole (DAPI)-supplemented antifade mounting medium for nuclear counterstaining. Micrographs were acquired using a ZEISS LSM 900 laser scanning confocal microscope or a ZEISS Axioscan 7 digital slide scanner (Carl Zeiss AG, Oberkochen, Germany). To evaluate pathological changes in myofiber size, the cross-sectional area (CSA) of individual fibers was measured using ImageJ software (National Institutes of Health, Bethesda, MD, USA).

### RNA extraction and reverse transcription quantitative PCR (RT-qPCR)

Total RNA was isolated from the tissue samples using the Direct-zol™ RNA Miniprep (R2050, Zymo Research, CA, USA) following the manufacturer’s protocols. Subsequently, cDNA was obtained using PrimeScript RT Master Mix (RR036A, Takara Bio, Shiga, Japan). For quantitative real-time PCR, the synthesized cDNA was mixed with target-specific primers and TB Green Premix (RR820A, Takara Bio, Shiga, Japan). The amplification reactions were conducted and analyzed on a qTOWER³ real-time PCR thermal cycler (Analytik Jena, Jena, Germany). *Gapdh* and *β-actin* were used as the endogenous reference genes for normalization. Relative mRNA expression levels were calculated using the standard 2^−ΔΔ*Ct*^ method. The following primers were used for this study: *Lipt1*-F (5’-TCA TCG GCA GGC ATC AGA ATC-3’), *Lipt1*-R (5’-CAT GTC GTG GTA CAC GGC TC −3’), *LplA*-F (5’-TGCCAAACTTCGCCGAAACC-3’), *LplA*-R (5’-CAGCAGATGCGAGAATGCCG-3’), *Gapdh*-F (5’-TCAAGAAGGTGGTGAAGCAG −3’), *Gapdh*-R (5’-GTTGAAGTCGCAGGAGACAA-3’), *β-actin*-F (5’-TGCTGTCCCTGTATGCCTCTG-3’)*, β-actin*-R (5’-TGATGTCACGCACGATTTCC-3’).

### RNA-sequencing

Total RNA was extracted from cardiac tissues using TRIzol reagent (Thermo Fisher Scientific, Waltham, MA, USA) following the manufacturer’s protocol. RNA concentration and purity were determined on a NanoDrop 2000 spectrophotometer (Thermo Fisher Scientific), and the integrity of each preparation was evaluated on an Agilent 5300 Bioanalyzer (Agilent Technologies, Santa Clara, CA, USA). Samples with a total yield above 1 µg, a concentration of at least 20 ng/µl and an RNA quality number (RQN) greater than 4.5 were carried forward to library preparation.

Strand-specific libraries were constructed with the Illumina Stranded mRNA Prep Ligation kit (Illumina, San Diego, CA, USA) from 1 µg of total RNA. Polyadenylated transcripts were captured on oligo(dT) magnetic beads and randomly fragmented in fragmentation buffer. First-strand cDNA was synthesized from random hexamer primers using SuperScript reverse transcriptase (Thermo Fisher Scientific), and dUTP was incorporated during second-strand synthesis to label the second strand. After the second strand had been digested with uracil-DNA glycosylase, the cDNA was end-repaired, phosphorylated and adenylated, and Y-shaped adaptors were ligated. The products were purified, size-selected for an average insert length of approximately 300 bp and enriched by PCR. Libraries were quantified with a Qubit 4.0 fluorometer (Thermo Fisher Scientific) and sequenced on a NovaSeq X Plus platform (Illumina) in 2 × 150 bp paired-end mode. Library preparation and sequencing were performed by Shanghai Majorbio Bio-pharm Biotechnology Co., Ltd. (Shanghai, China).

Raw reads were processed with fastp to remove adaptor sequences and low-quality bases, and reads containing more than 10% ambiguous bases or shorter than 20 bp after trimming were discarded. Clean reads were aligned to the mouse reference genome (GRCm39) with HISAT2, and transcript assembly and abundance estimation were performed with StringTie and RSEM, respectively, with expression levels expressed as transcripts per million (TPM). Principal component analysis was carried out on the normalized expression matrix, and differentially expressed genes (DEGs) were identified with DESeq2 using a threshold of absolute log2 fold change of 1 or greater and a Benjamini-Hochberg-adjusted P value below 0.05. Volcano plots were generated to display the distribution of DEGs. Gene Ontology (GO) and Kyoto Encyclopedia of Genes and Genomes (KEGG) enrichment analyses of the DEGs were performed with Goatools and the Python SciPy package, respectively, and terms with a Bonferroni-corrected P value below 0.05 were considered significantly enriched. Bioinformatic analyses were conducted on the online Majorbio Cloud Platform (https://cloud.majorbio.com).

The complete RNA sequencing dataset was deposited in the NCBI BioProject database under accession number PRJNA1550529.

### Western blot

Tissue samples (approximately 20 mg) were finely minced and homogenized in 200 µL of radioimmunoprecipitation assay (RIPA) lysis buffer (P0013B, Beyotime Biotechnology, Shanghai, China) containing 1 mM phenylmethylsulfonyl fluoride (PMSF) using a refrigerated high-throughput tissue homogenizer (SCIENTZ-48LT, Ningbo Scientz Biotechnology, China) for 5 min. Following a 30-min incubation on ice to ensure thorough lysis, the mixtures were centrifuged at 12,000 × g for 30 min at 4 °C, and the supernatants were carefully collected. Total protein concentrations were measured using a Bradford protein assay kit (P0006, Beyotime Biotechnology, Shanghai, China) according to the manufacturer’s instructions. Protein samples were mixed with sodium dodecyl sulfate-polyacrylamide gel electrophoresis (SDS-PAGE) loading buffer (Takara Bio, Japan) and heated at 98 °C for 10 min. Proteins were loaded on SDS-PAGE gels and transferred onto nitrocellulose (NC). The NC membranes were blocked using a protein-free rapid blocking buffer (PS108P, Epizyme Biotech, Shanghai, China) for 10 min at room temperature, and incubated overnight at 4 °C with the following primary antibodies: anti-LplA antibody (CUSBIO, CSB-PA510777HA01ENU, China, 1:1000), anti-lipoic acid antibody (Abcam, ab58724, UK, 1:2000), anti-GAPDH antibody (Proteintech, 60004, USA, 1:5000), anti-TOMM20 antibody (Abcam, ab78547, UK, 1:1000), OXPHOS cocktail (Abcam, ab110413, UK, 1:1000), anti-pAkt antibody (CST, 4060S, USA, 1:1000), anti-Akt antibody (CST, 9272S, USA, 1:1000), anti-LC3 antibody (1:1000), and anti-β-actin (Abclonal, AC026, China, 1:1000). Anti-rabbit immunoglobulin G (IgG) (LI-COR, 926-32211, USA, 1:10,000) and anti-mouse IgG (LI-COR, 926-68070, USA, 1:10,000) were applied as secondary antibodies. ChemiDoc MP Imaging System (12003154, Bio-Rad Laboratories, CA, USA) was used for imaging, and densitometric quantification was performed with ImageLab software (Bio-Rad Laboratories, CA, USA).

### Gene and protein nomenclature

Human genes are italicized in all caps; mouse gene names are italicized with only the first letter capitalized. Proteins for both humans and animals are in all caps without italics.

### Statistical analysis

Data are presented as mean ± SD. All analyses were performed using GraphPad Prism 10 software (GraphPad Software Inc., La Jolla, CA). Survival curves were generated using the Kaplan-Meier method and compared by the log-rank (Mantel-Cox) test. Two-group comparisons were performed using unpaired two-tailed t tests. Comparisons among three or more groups were performed using one-way analysis of variance (ANOVA) followed by Tukey’s multiple comparisons test. Statistical significance was defined as \**P* < 0.05, \*\**P* < 0.01, \*\*\**P* < 0.001, and \*\*\*\**P* < 0.0001.

## DATA AVAILABILITY STATEMENT

All data are available in the main text or the supplementary materials. Further information and requests for resources and reagents should be directed to and will be fulfilled by the corresponding author.

## ACKNOWLEDGEMENTS

We thank the staff from the Laboratory Animal Resources Center and the Advanced Biomedical Technology Core Facility at Westlake University for advice and assistance in data collection and discussion. This work was financially supported by the National Key R&D Program of China (2022YFA0912000), Zhejiang Key Laboratory of Low-Carbon Intelligent Synthetic Biology (2024ZY01025), Zhejiang Key Laboratories Project (2024E10053), "Pioneer" and "Leading Goose" R&D Program of Zhejiang (2025C01094).

## AUTHOR CONTRIBUTIONS

Conceptualization: JY, ZH, AZ

Methodology: JY

Investigation: JY, ZL

Visualization: JY, ZL

Funding acquisition: AZ

Project administration: JY, ZH, AZ

Supervision: ZH, AZ

Writing-original draft: JY

Writing-review & editing: JY, ZH, RY, AZ

