## Supplemental figures for "LplA-mediated rescue of lethal myopathy and cardiomyopathy in mitochondrial lipoylation-deficient mammalian caused by *Lipt1*"

Junru YU *et al.*

**The PDF file includes:**

Figure S1. Characterization of Δ*Lipt1* and *Lipt1*-cKO mice.

Figure S2. Hematological and plasma biochemical analyses.

Figure S3. Systemic physical status and skeletal muscle architecture analyses.

Figure S4. Cardiac transcriptomic analysis.

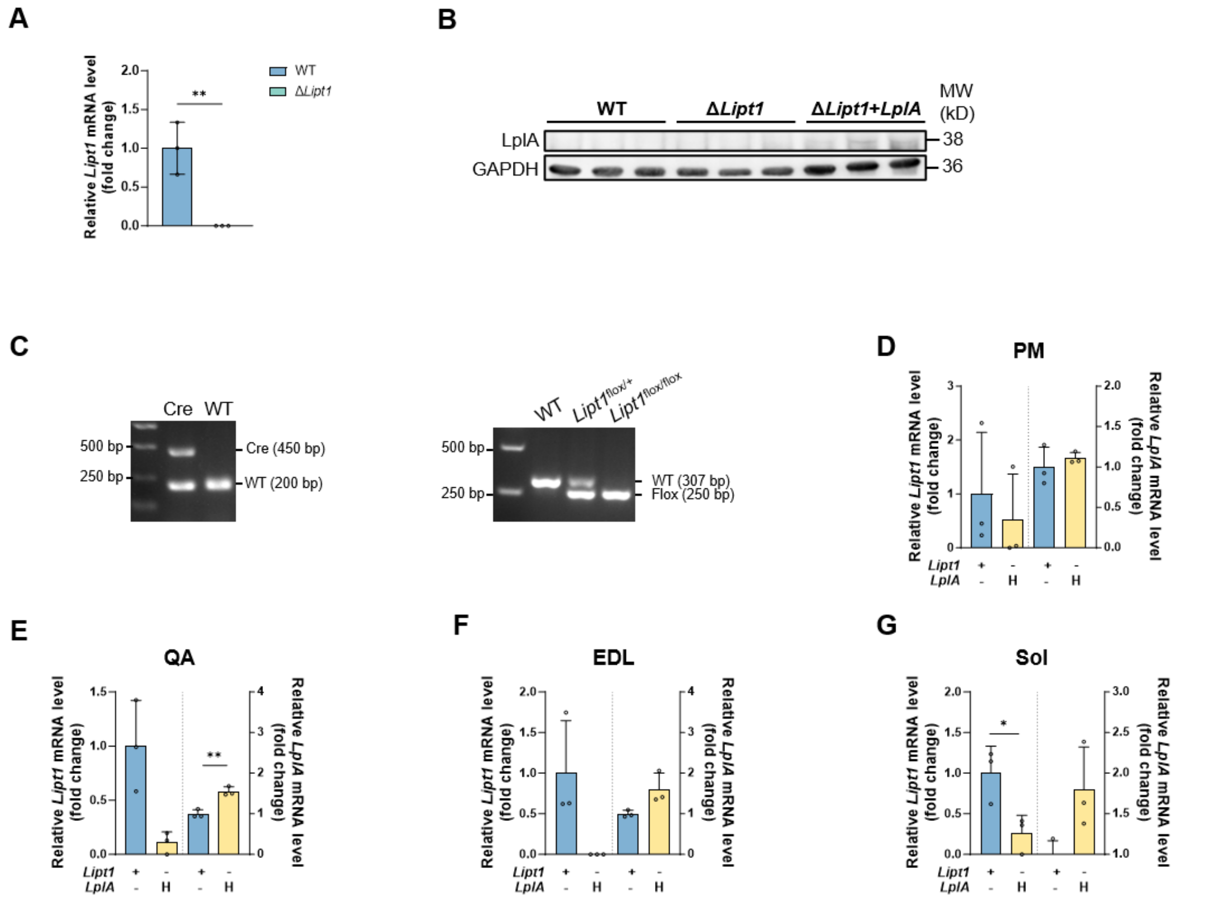


**Figure S1. Characterization of Δ*Lipt1* and *Lipt1*-cKO mice.**

**(A)** Relative *Lipt1* mRNA levels in WT and Δ*Lipt1*, determined by RT-qPCR. *Actin* was used as the internal control for normalization

**(B)** Immunoblots of LplA in WT, Δ*Lipt1* and Δ*Lipt1*+*LplA* mice. GAPDH was used as the control.

**(C)** Representative genotyping results of WT, *Ckmm*-Cre, *Lipt1*$^{\text{flox/+}}$, *Lipt1*$^{\text{flox/flox}}$ mice.

**(D-G)** Relative *Lipt1* (left axis) and *LplA* (right axis) mRNA levels in PM (D), QA (E), EDL (F) and Sol (G) of wild-type (WT) and *Lipt1*-cKO+LplA(H)+LA mice, determined by RT-qPCR. *GAPDH* was used as the internal control for normalization

n = 3 mice per group. Data are presented as mean ± SD. Statistical significance was determined by unpaired two-tailed t-test. *$P < 0.05$.

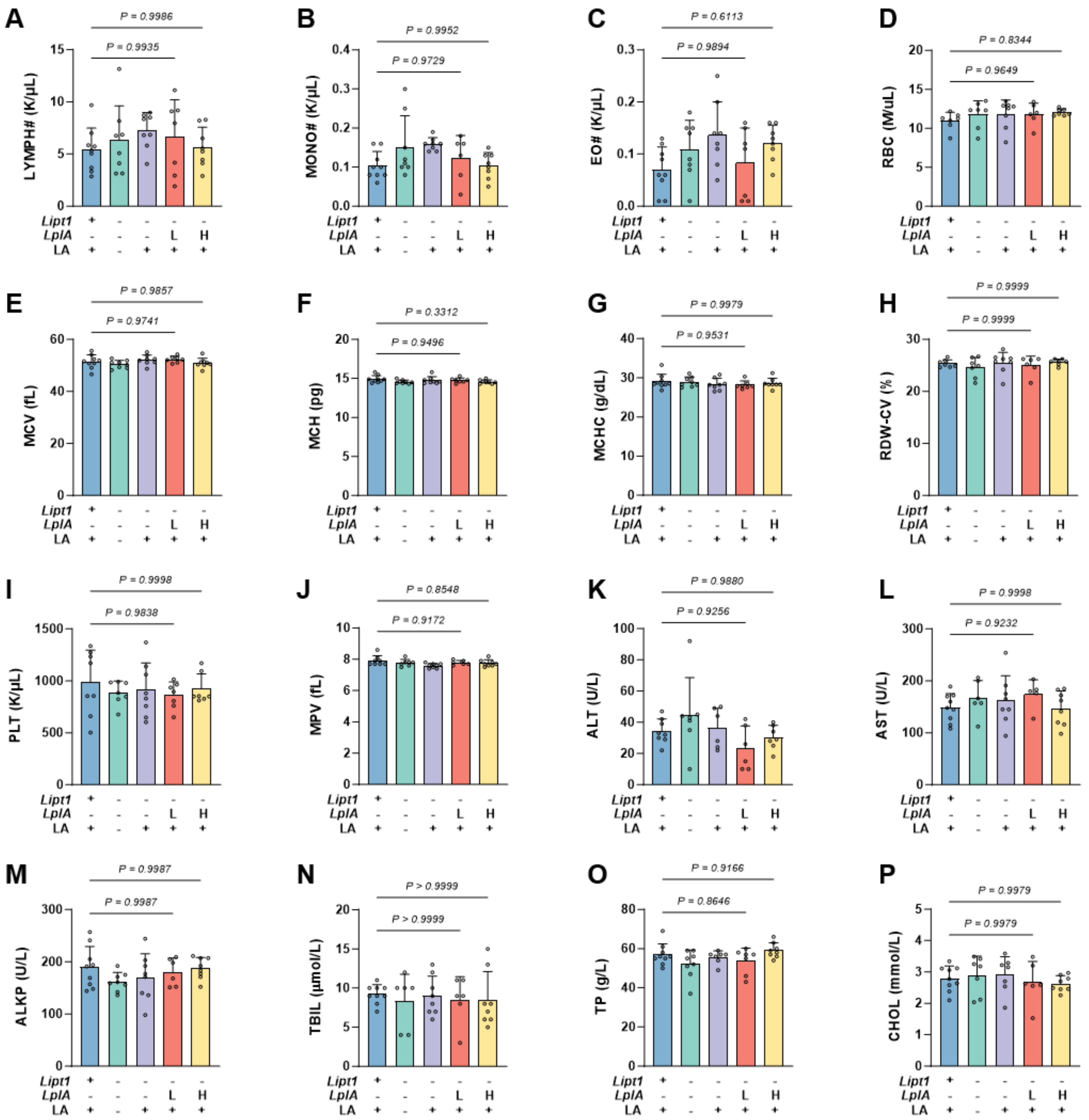

**Figure S2. Hematological and plasma biochemical analyses.**

**(A-J)** Complete blood count parameters in the Control+LA, *Lipt1*-cKO, *Lipt1*-cKO+LA, *Lipt1*-cKO+*LplA*(L)+LA and *Lipt1*-cKO+*LplA*(H)+LA mice. (A) Lymphocyte count (LYMPH#). (B) Monocyte count (MONO#). (C) Eosinophil count (EO#). (D) Red blood cell count (RBC). (E) Mean corpuscular volume (MCV). (F) Mean corpuscular hemoglobin (MCH). (G) Mean corpuscular hemoglobin concentration (MCHC). (H) Red cell distribution width (RDW-CV). (I) Platelet count (PLT). (J) Mean platelet volume (MPV).

**(K-P)** Plasma biochemistry parameters in the indicated groups. (K) Alanine aminotransferase (ALT). (L) Aspartate aminotransferase (AST). (M) Alkaline phosphatase (ALKP). (N) Total bilirubin (TBIL). (O) Total protein (TP). (P) Cholesterol (CHOL).

n = 5-9 mice per group. Data are presented as mean ± SD. Statistical significance was determined by one-way ANOVA followed by Tukey’s multiple comparisons test. No significant differences were detected between the indicated groups.

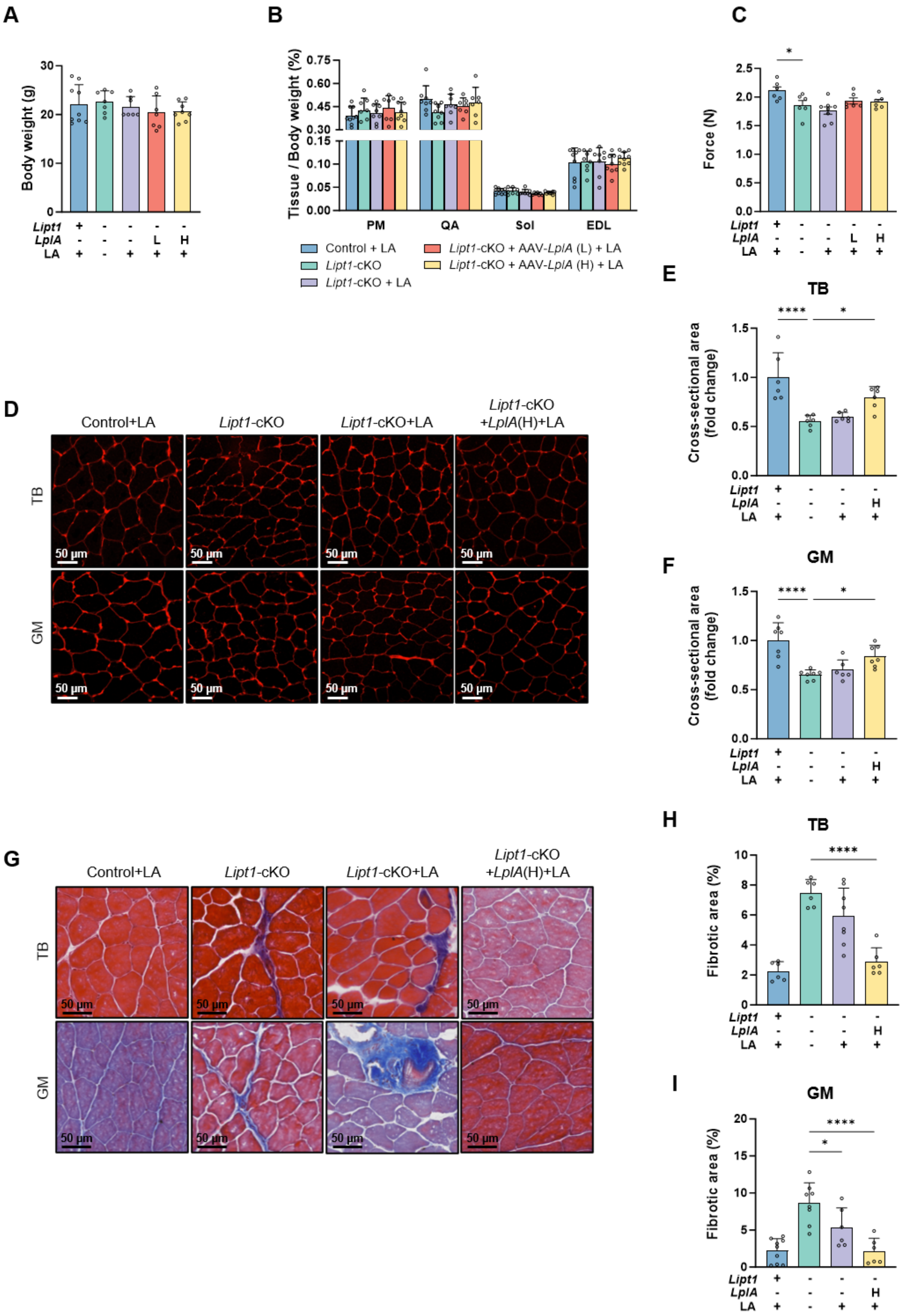


**Figure S3. Systemic physical status and skeletal muscle architecture analyses.**

**(A)** Body weights of Control+LA, *Lipt1*-cKO, *Lipt1*-cKO+LA, *Lipt1*-cKO+*LplA*(L)+LA and *Lipt1*-cKO+*LplA*(H)+LA mice.

**(B)** Weights of PM, QA, Sol and EDL normalized to body weight.

**(C)** Maximal grip force measured by grip strength test.

**(D)** Representative WGA-stained cross-sections of TB (top) and GM (bottom) muscles.

**(E and F)** Myofiber cross-sectional area in TB (E) and GM (F), expressed relative to Control+LA.

**(G)** Representative Masson's trichrome-stained TB and GM muscles.

**(H and I)** Interstitial fibrotic area in TB (H) and GM (I).

Scale bars, 50 μm (D and G). n = 6-9 mice per group. Data are presented as mean ± SD. Statistical significance was determined by one-way ANOVA followed by Tukey's multiple comparisons test. *$P < 0.05$, **$P < 0.01$ and ****$P < 0.0001$.

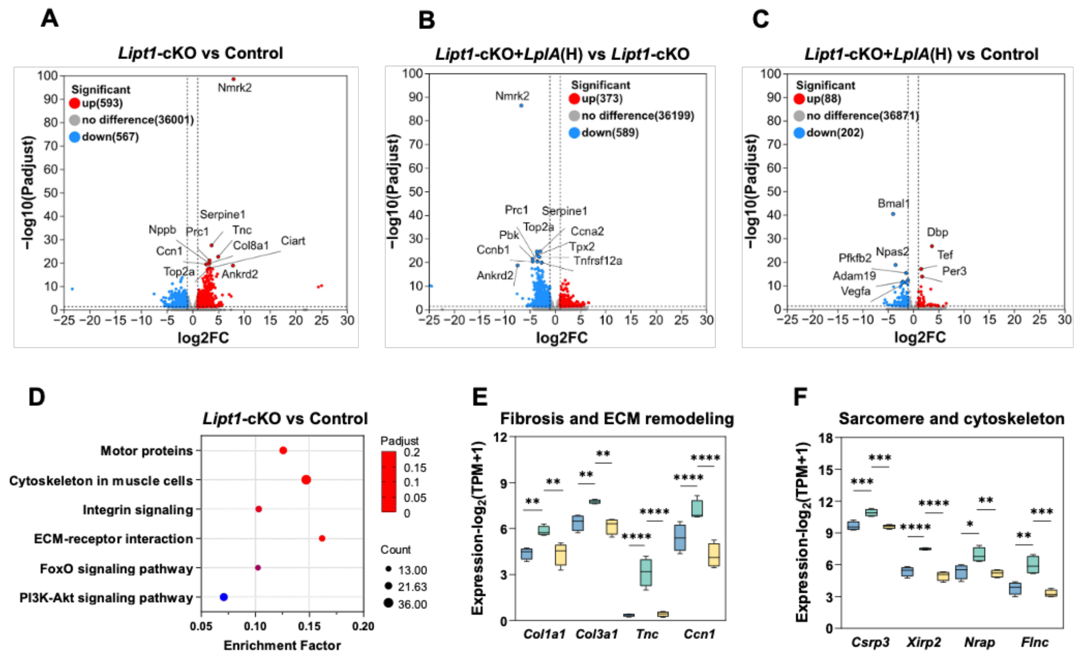


**Figure S4. Cardiac transcriptomic analysis.**

**(A-C)** Volcano plots of differentially expressed genes for *Lipt1*-cKO versus Control+LA hearts (A), *Lipt1*-cKO+LplA(H) versus *Lipt1*-cKO hearts (B) and *Lipt1*-cKO+LplA(H) versus Control+LA hearts (C). Red and blue indicate significantly upregulated and downregulated genes, respectively; grey indicates genes that did not pass the significance threshold. Selected genes are labeled.

**(D)** KEGG pathway enrichment of differentially expressed genes between *Lipt1*-cKO and Control+LA hearts. The color scale represents the adjusted P value, and the dot size represents the number of genes.

**(E)** Expression of fibrosis and ECM remodeling genes (*Col1a1*, *Col3a1*, *Tnc* and *Ccn1*). Transcript abundance is shown as log2(TPM+1), where TPM denotes transcripts per million.

**(F)** Expression of sarcomere and cytoskeleton genes (*Csrp3*, *Xirp2*, *Nrap* and *Flnc*).

n = 4 mice per group. Data are presented as mean ± SD. Statistical significance was determined by one-way ANOVA followed by Tukey's multiple comparisons test. *$P < 0.05$, **$P < 0.01$ and ***$P < 0.001$.
